# Beyond Global Contrasts: Paired-Unit Activation Steering for Context-Aware Protein Sequence Design

**DOI:** 10.64898/2026.11.09.765362

**Authors:** Yingtong Zhou, Xiaoning Qian, Byung-Jun Yoon

**Affiliations:** Department of Electrical and Computer Engineering, Texas A&M University, TX 77843, USA; Department of Computer Science and Engineering, Texas A&M University, TX 77843, USA; Computing and Data Sciences, Brookhaven National Laboratory, Upton, NY 11973, USA

**Keywords:** Protein Language Models, Paired-Unit Activation Steering, Protein Optimization, Context-Aware Protein Design

## Abstract

Activation steering in protein language models enables controllable sequence generation without changing the model weights. However, existing methods typically construct steering vectors from global differences between property-positive and property-negative protein populations, overlooking how property-improving directions vary across different protein sequence backgrounds. To address this, we propose *Paired-Unit local steering of embeddings (PULSE)*, which matches positive- and negative-property proteins by maximizing their representation cosine similarity and constructs steering vectors by aggregating similarity-weighted, unit-normalized differences between matched pairs. Across three protein optimization objectives–solubility, immunogenicity, and toxicity– PULSE consistently outperforms global population-centroid steering in both final predicted property scores and optimization success rates. Further analysis reveals that conventional static steering-vector diagnostics cannot adequately explain optimization performance. To characterize steering reliability during iterative sequence editing, we introduce *Local Directional Support (LDS)*, which measures how well successive representation changes align with property-improving directions from similar protein backgrounds. We show that LDS captures differences in trajectory reliability and enables predictor-free candidate prioritization, enriching successful protein designs without requiring property predictions during candidate selection. Together, these findings demonstrate the importance of context-aware steering and trajectory-level evaluation for controllable protein sequence optimization.

## 1 Introduction

Protein sequence optimization aims to improve desirable properties while preserving native structural and functional characteristics. Recent advances in pretrained protein language models (pLMs) have facilitated this process by learning representations that capture evolutionary, structural, and functional information from large-scale sequence datasets [18,11,13]. These models have shown promising capabilities in mutation-effect prediction and protein sequence generation [14,6]. However, guiding pLMs toward specific properties remains challenging, especially when labeled data are limited or fine-tuning is computationally expensive.

Activation steering offers a lightweight alternative by controlling model behavior through modifications to internal representations without updating model parameters [21,17]. It constructs steering directions from contrasting examples and applies them to guide generation toward desired properties. Building on this idea, Activation Steering based Protein Optimization (ASPO) [7] computes a steering vector from the difference between the mean representations of property-positive and property-negative proteins, using it to guide iterative residue-level sequence optimization. Recent work has also explored uncertainty-aware steering to improve optimization reliability [15].

Despite its effectiveness, population-level steering relies on a global contrast that may overlook differences among individual protein backgrounds. Protein fitness landscapes are often characterized by epistasis, where the effect of a mutation depends on other residues in the sequence [22,19,20]. Consequently, changes that improve a property in one protein background may not provide the same benefit in another. This suggests that property-associated directions in learned representation space may also vary across sequence backgrounds. By averaging representations across heterogeneous proteins, global centroid-based steering as in ASPO [7] may obscure these local differences and produce directions that are less suitable for individual starting sequences.

To address this limitation, we propose **Paired-Unit Local Steering of Embeddings (PULSE)**, a context-aware approach to constructing property-associated steering vectors. Rather than directly contrasting population centroids, PULSE matches property-positive and property-negative proteins according to their representation similarity. It then aggregates similarity-weighted, unit-normalized differences between matched pairs to construct the steering direction. This design aims to preserve information from comparable protein backgrounds while reducing the influence of differences in pairwise displacement magnitudes.

We evaluate PULSE against conventional centroid-based steering across three protein optimization objectives: solubility, immunogenicity, and toxicity. Beyond improving optimization performance, we investigate how steering directions relate to the reliability of iterative sequence editing. Conventional diagnostics based on static steeringvector properties do not necessarily reflect whether individual optimization steps follow property-improving directions. To examine this behavior, we introduce **Local Directional Support (LDS)**, a trajectory-level metric that evaluates the alignment between successive representation changes and property-associated directions from similar protein backgrounds. Our experiments show that LDS captures differences in optimization reliability and supports predictor-free candidate prioritization.

Our main contributions are summarized as follows:

1. We demonstrate that property-associated directions in protein representation space exhibit systematic background dependence, motivating context-aware steering.
2. We propose **PULSE** to construct steering vectors from similarity-matched, unitnormalized protein representation differences, which improves optimization outcomes across three protein properties.
3. We show that static steering-vector diagnostics are insufficient to explain optimization performance and introduce **LDS** to characterize the reliability of iterative sequence-editing trajectories.
4. We demonstrate that LDS enables predictor-free prioritization of optimized protein candidates, enriching successful designs for downstream experimental screening.

## 2 Methods

### 2.1 Protein Sequence Optimization with Activation Steering

For sequence *x*, let 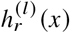 be the representation of residue *r* at layer *l*, (*h*)^(*l*)^ *x* its mean over residues, and *v*^(*l*)^ a property-specific steering vector. Following ASPO [7], each step ranks residues of *x*^*t*^ by 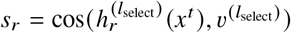 and masks the lowest-scoring sites. Here *l*_select_ denotes the backbone layer used for site selection.

During decoding, the backbone steers each hidden state while preserving its original norm:

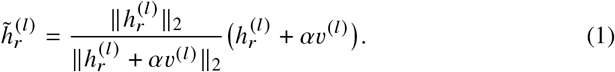

The decoded sequence becomes the optimized *x*^*t*+1^. PULSE adpots the same site selection and decoding as ASPO; only the construction of *v*^(*l*)^ differs, as described next.

### 2.2 Steering Vector Construction

#### ASPO baseline

ASPO constructs a global steering vector from the difference between the mean representations of property-positive and property-negative proteins:

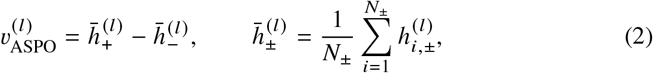

where *N*_±_ counts the number of sequences in each class. Thus, this global contrast does not preserve individual correspondences.

#### Paired-Unit Local Steering of Embeddings (PULSE)

PULSE instead constructs steering directions from matched positive–negative pairs (Fig. 1). At the pairing layer *l*_pair_, we compute

**Fig. 1:**
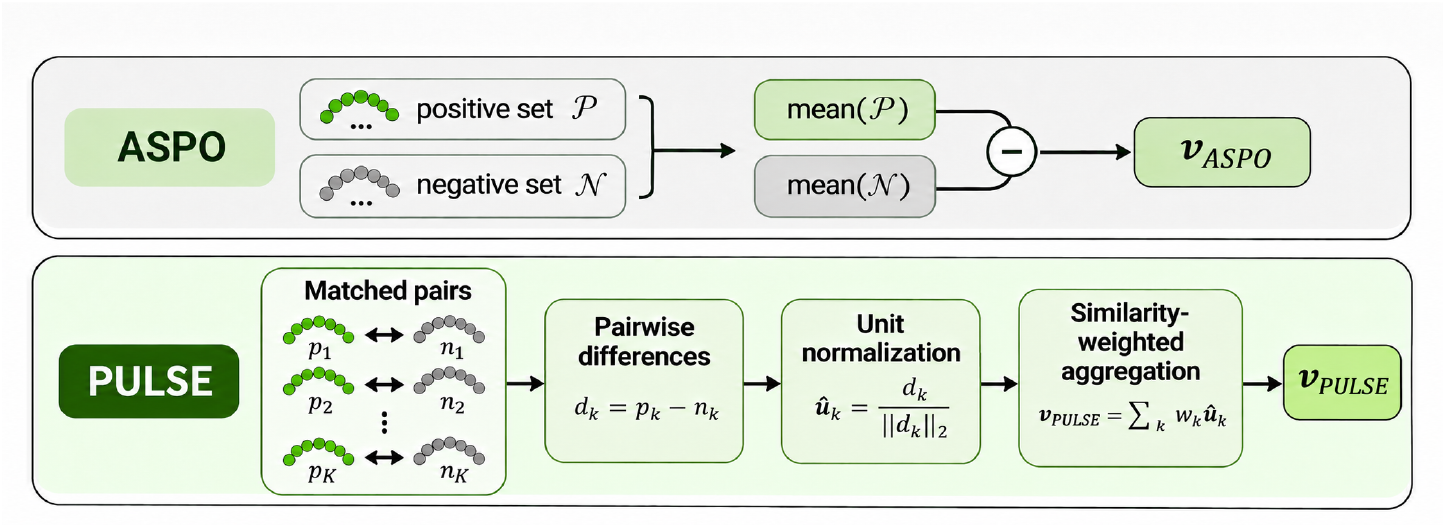
ASPO and PULSE steering-vector construction. ASPO uses the difference between global mean representations, whereas PULSE constructs steering vectors from similarity-matched positive–negative pairs.

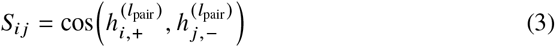

and use the Hungarian algorithm [9] to obtain *K* = min (*N*_+_, *N*_−_) disjoint pairs that maximize total similarity. Let *i*_*k*_, *j*_*k*_ denote matched pair *k* and 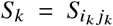. This pairing is computed once and reused to construct steering vectors at every layer *l*.

For each matched pair, the representation difference at layer *l* is

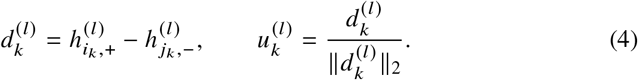

We aggregate these unit-normalized directions using similarity-based softmax weights:

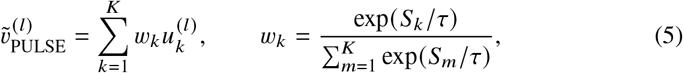

where *τ* = 0.10. The pairing-layer scores *S*_*k*_, and therefore the weights *w*_*k*_, are shared across layers and do not depend on the optimization target.

To ensure a fair comparison, the resulting vector is rescaled to match the norm of the ASPO steering vector. Thus, PULSE differs from ASPO in steering-direction while retaining the same steering magnitude and downstream optimization procedure.

For either method, let *x*^0^ be a target sequence and *x*^*T*^ its sequence after *T* optimization steps. An external property predictor *f* provides the final score *f* (*x*^*T*^) and the retrospective success label

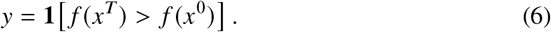

The predictor is used only for evaluation, not to guide sequence editing.

### 2.3 Steering Reliability Diagnostics

We first evaluated static steering-vector reliability using global reproducibility and Local Static Vector Support (LSVS). Global reproducibility was assessed through *B* = 200 bootstrap reconstructions and their angular deviations from the full-data steering vector [3]. LSVS measures the fraction of property-associated directions from similar protein backgrounds that align positively with the static steering vector. Detailed formulations are provided in Appendix A.2.

To evaluate reliability during iterative optimization, we introduce **Local Directional Support (LDS)**. For each optimization step *t* → *t* + 1, we compute the realized representation displacement 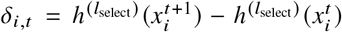. From an independent reference pool, we retrieve the *K* = 32 property-improving directions whose pair backgrounds are most similar to the starting protein 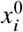 at *l*_pair_. Each pair background is the normalized midpoint of the individually normalized positive and negative protein representations. This neighborhood is fixed throughout each trajectory, so PULSE and ASPO are evaluated against the same starting-context reference for each target. For a nonzero displacement, step-level LDS is defined as

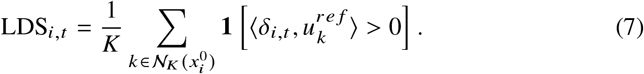

Here, 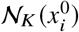 denotes the selected local reference directions and 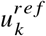 denotes a unit direction from the reference pool; a zero displacement receives LDS 0.5. Trajectory-level reliability is summarized by the lower quartile of step-level LDS values:

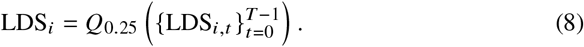

This aggregation emphasizes weakly supported steps rather than average trajectory alignment.

Finally, candidates were ranked by trajectory-level LDS without using property predictions during selection. Ranking quality was evaluated retrospectively using predictor labels, AUROC, and success enrichment among the top-ranked 25% of candidates.

### 2.4 Data Availability

The implementation and usage instructions are available at https://github.com/TeresaZhouTamu/Paired_Unit_Steering.

## 3 Results

### 3.1 Evaluation tasks and settings

We evaluated ASPO and PULSE on solubility, immunogenicity, and toxicity optimization using independent test sets of 100, 100, and 50 target proteins, respectively. Both methods used ESMC-600M and the same optimization settings: *T* = 4 steps, 2 smutation sites per step, and deterministic decoding (temperature = 0). The (*l*_select_, *l*_pair_, *α*)settings were specified in Appendix C. External property predictors scored the original and final sequences retrospectively. Dataset and predictor details are given in Appendix B.

### 3.2 Paired-Unit steering improves controllable protein sequence optimization

We first evaluated whether PULSE improves protein sequence optimization compared with ASPO. Across solubility, immunogenicity, and toxicity, PULSE achieved higher mean final predicted property scores than ASPO (Fig. 2a), with the largest improvement observed for toxicity. Paired comparisons confirmed statistically significant improvements across all three objectives after multiple-testing correction (Appendix A.1, Table 2).

**Fig. 2:**
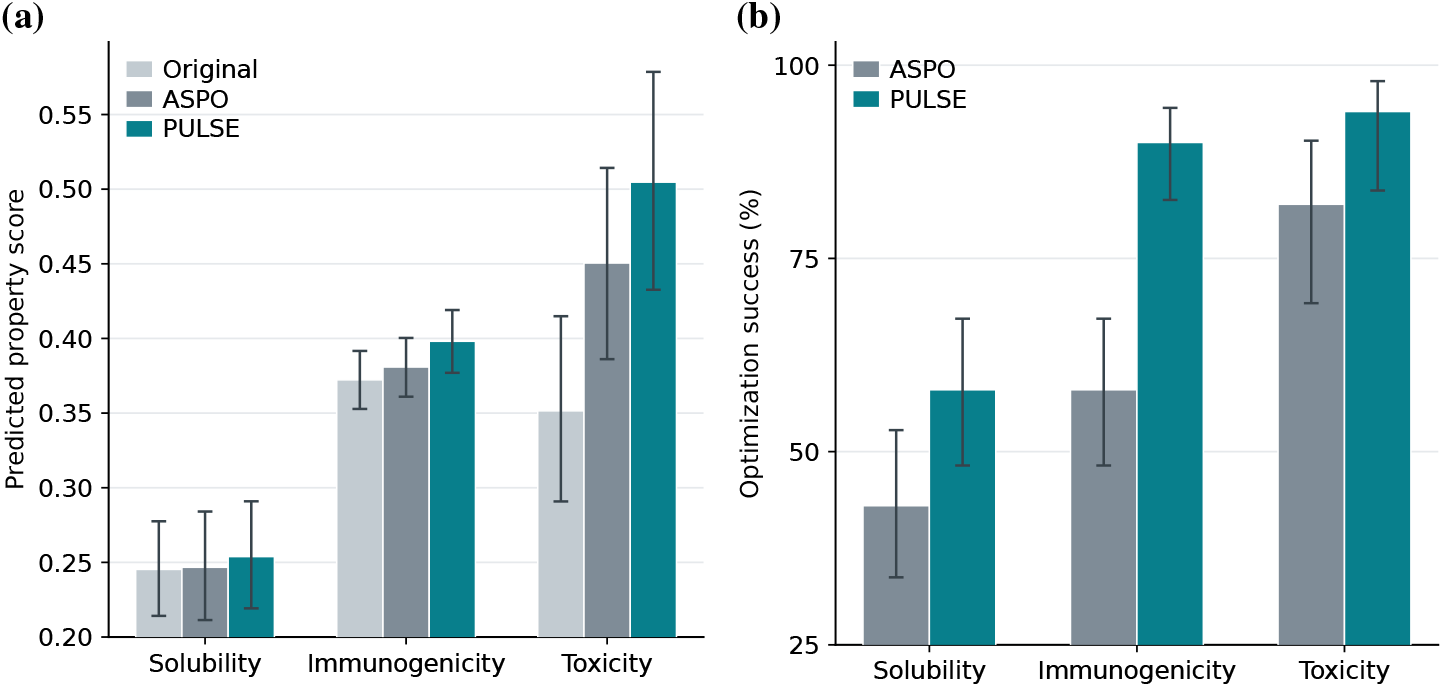
PULSE improves predicted optimization outcomes. (a) Mean property scores before and after optimization with 95% bootstrap CIs. (b) Optimization success rates with 95% Wilson CIs.

We further examined optimization success, defined as the proportion of targets whose final predicted scores exceeded their original scores. PULSE achieved higher success rates across all three properties (Fig. 2b), with the largest increase observed for immunogenicity. Together, these results demonstrate that PULSE improves both final predicted property scores and the proportion of successfully optimized targets. Because the optimization and decoding procedures were shared, these results support the contribution of PULSE steering-direction construction to the observed performance gain.

### 3.3 Property-improving directions exhibit systematic background dependence

Having established PULSE’s performance advantage, we next examined whether the pair-specific directions retained during its construction capture variation across protein backgrounds. If these contrasts contain context-dependent information, pairs with similar backgrounds should exhibit more closely aligned property-associated directions.

For each matched positive–negative pair, we defined its background as the normalized midpoint of the two unit-normalized sequence representations and its property-associated direction as the normalized negative-to-positive representation difference. We then compared background cosine similarity with direction cosine similarity across pairs. Across all three properties, more similar protein backgrounds tended to exhibit more similar property-associated directions (Fig. 3). The positive Spearman correlations support this global trend across all three properties, with the strongest association observed for toxicity (Table 1).

**Table 1.** Background dependence and local coherence.

| Property | *ρ* | Local coherence |  |
|---|---|---|---|
|  |  | Observed | Null |
| Solubility | 0.163 | 0.217 | 0.084 |
| Immunogenicity | 0.316 | 0.258 | 0.081 |
| Toxicity | 0.568 | 0.302 | 0.161 |

**Property:** optimization objective.
***ρ***: Spearman correlation.
**Observed:** local directional coherence.
**Null:** permutation-mean coherence.

**Table 2.** Mean final predicted scores and paired PULSE–ASPO differences.

| **Property** | **ASPO** | **PULSE** | **Δ** | **95% CI** | **Holm *p*** |
|---|---|---|---|---|---|
| Solubility | 0.24671 | 0.25386 | 0.00715 | [0.00151, 0.01353] | 0.00240 |
| Immunogenicity | 0.38084 | 0.39808 | 0.01724 | [0.01172, 0.02282] | < 10<sup>−5</sup> |
| Toxicity | 0.45051 | 0.50469 | 0.05417 | [0.02447, 0.08680] | 0.00019 |

**Table 3.**
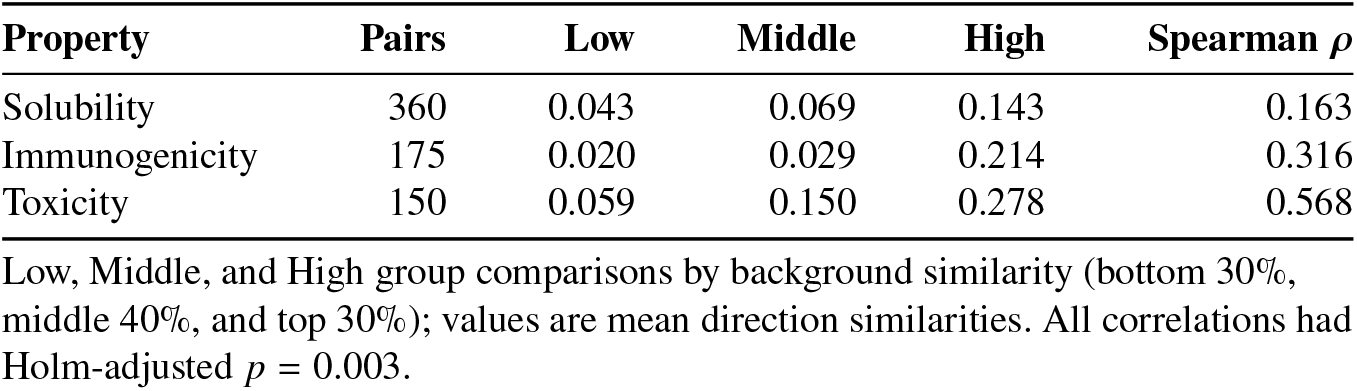
Background similarity and property-direction agreement.

**Table 4.** Summary of dataset configurations and sequence counts after cleaning.

| **Property** | **Dataset Role** | **Raw *N*** | **Cleaned *N* (Pos / Neg)** |
|---|---|---|---|
| Solubility | Main Construction | 720 | 360 / 360 |
|  | Test Target Set | 100 | 0 / 100 |
|  | LDS Reference Pool | 400 | 200 / 200 |
| Immunogenicity | Main Construction | 394 | 175 / 175 |
|  | Test Target Set | 100 | 0 / 100 |
|  | LDS Reference Pool | 250 | 125 / 125 |
| Toxicity | Main Construction | 300 | 150 / 150 |
|  | Test Target Set | 50 | 0 / 100 |
|  | LDS Reference Pool | 250 | 125 / 125 |

**Fig. 3:**
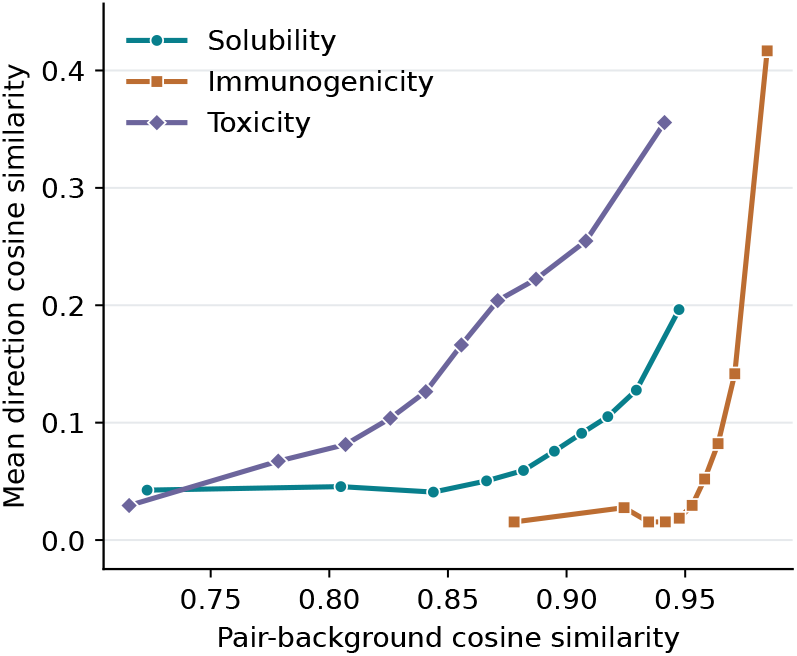
Similar protein backgrounds are associated with more aligned directions. Each curve shows average directional agreement across different levels of background similarity.

To complement this global association, we also examined directional coherence within neighborhoods of similar pair backgrounds. Neighboring backgrounds exhibited greater directional agreement than expected under randomized background–direction assignments. Both the global association and local coherence were statistically significant across all three properties (Table 1).

Rather than random deviations around a global vector, paired property directions exhibit a systematic structure tied to the protein background supporting PULSE’s use of retaining pair-specific information prior to aggregation. While providing an empirical rationale for PULSE’s performance advantage over ASPO, this observation also raises a further question: does this context-aware advantage also manifest as greater steering reliability?

### 3.4 Static steering-vector uncertainty does not capture optimization reliability

To investigate whether static steering-vector reliability explains PULSE’s optimization advantage, we examined two complementary diagnostics: global reproducibility and target-conditioned Local Static Vector Support (LSVS) (Appendix A.2). The former measures the stability of steering directions, while the latter evaluates their alignment with property-improving directions from similar protein backgrounds.

Global reproducibility was assessed through bootstrap-based angular deviations from the full-data steering vector [3]. Despite achieving better optimization outcomes, PULSE exhibited greater mean angular deviation than ASPO across all three properties, particularly toxicity (Fig. 4a). Meanwhile, LSVS showed no statistically significant differences between the two methods across any of the three properties (Fig. 4b).

**Fig. 4:**
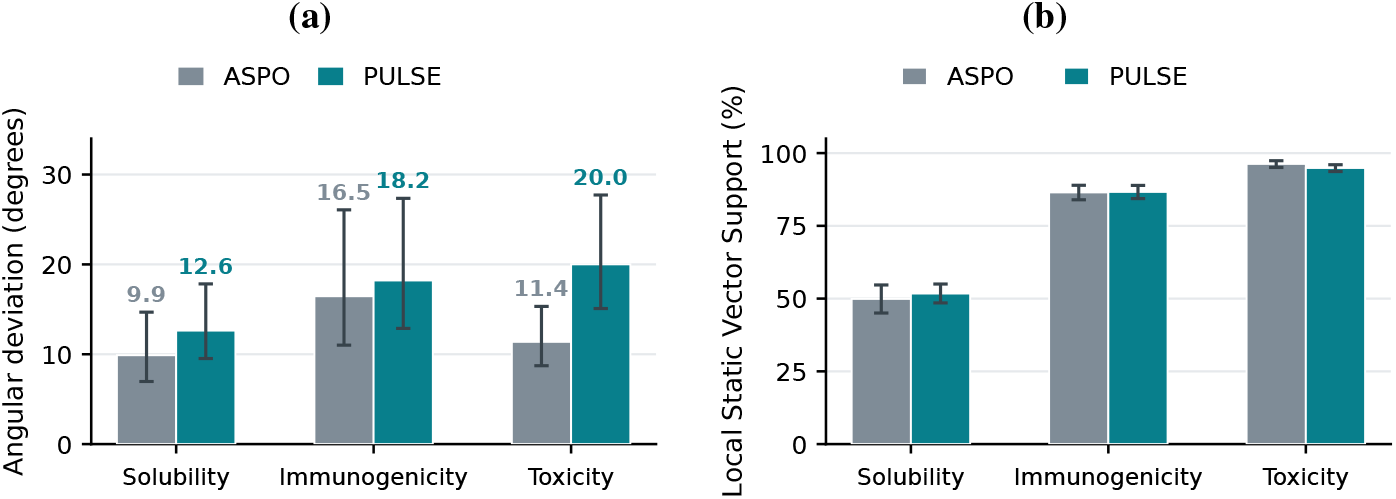
Static-vector reliability does not explain PULSE’s gains. (a) Mean bootstrap angular deviation; whiskers show the central 95% range. (b) Mean Local Static Vector Support (LSVS), the fraction of nearby property directions aligned with the static vector.

These results indicate that neither global reproducibility nor static local compatibility explains PULSE’s optimization advantage. While both assessments characterize the static, intended intervention direction, the realized sequence design also depends on the specific mutations selected during iterative decoding. We therefore extend our local evaluation from static vectors to the realized optimization trajectory, asking whether its intermediate representation changes are supported by property-improving directions from similar protein contexts.

### 3.5 Local Directional Support quantifies reliability along the realized optimization trajectory

Since static steering-vector diagnostics failed to explain PULSE’s optimization advantage, we shifted our focus to the *realized optimization trajectory*. While endpoint metrics capture final property improvements, they do not reveal whether intermediate editing steps follow directions supported by similar protein backgrounds. We therefore used **Local Directional Support (LDS)** to evaluate the alignment between realized representation shifts and locally supported property-improving directions, as defined in Section 2.3.

Across all three properties, PULSE trajectories exhibited consistently higher LDS than ASPO, with paired 95% confidence intervals for the mean differences excluding zero (Fig. 5). This indicates that PULSE’s editing steps were more consistently aligned with property-improving directions from comparable protein contexts.

**Fig. 5:**
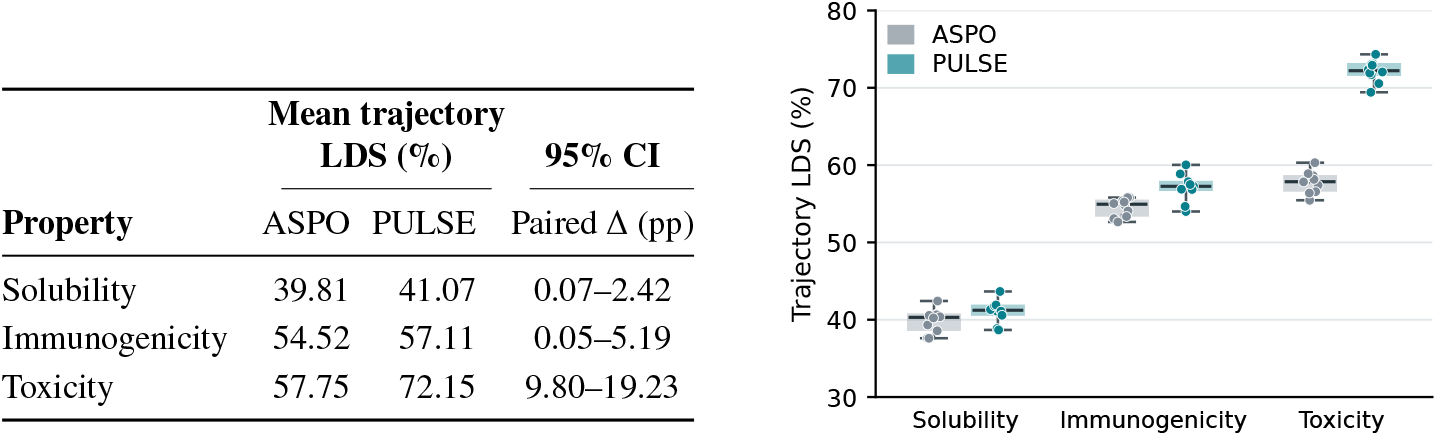
Local support along optimization trajectories. The table reports mean LDS and 95% paired confidence intervals. The plot shows ten reference pools; boxes mark medians and interquartile ranges, whiskers the observed range, and dots individual pools.

We further examined sensitivity to reference-set composition by evaluating LDS across 10 independently sampled reference pools, held out from steering-vector construction. PULSE maintained higher mean LDS than ASPO across all three properties, demonstrating that the observed advantage was robust to reference selection (Fig. 5).

Together, these findings suggest that trajectory-level directional support provides a more informative characterization of PULSE’s optimization behavior than static steering-vector diagnostics. Although PULSE’s steering vectors were not consistently more stable, its realized optimization trajectories showed stronger alignment with context-matched property-improving directions.

### 3.6 LDS enables reliability-guided prioritization of optimized proteins

Having established that LDS quantifies trajectory-level alignment with context-matched property trends, we next evaluated its utility for prospectively prioritizing candidate sequences for experimental testing. In practical protein engineering, wet-lab validation is constrained by throughput and cost, making candidate selection critical [16,2]. This is especially important when property measurements are too limited to train a reliable predictor for screening [1,12]. We hypothesized that trajectory-level LDS could serve as a general, oracle-free reliability filter to rank promising designs prior to synthesis.

To evaluate candidate prioritization without relying on a property model during selection, we ranked optimized sequences strictly in descending order of their LDS scores, using an external validator *f* (·) solely to establish retrospective ground-truth success labels 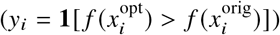. As shown in Fig. 6b, filtering candidates by LDS produces a consistently descending success profile across both ASPO and PULSE. This decay confirms that LDS functions as an effective, method-agnostic prioritization signal: across different steering paradigms, sequences traversed along trajectories with higher local support are systematically more likely to yield successful edits.

**Fig. 6:**
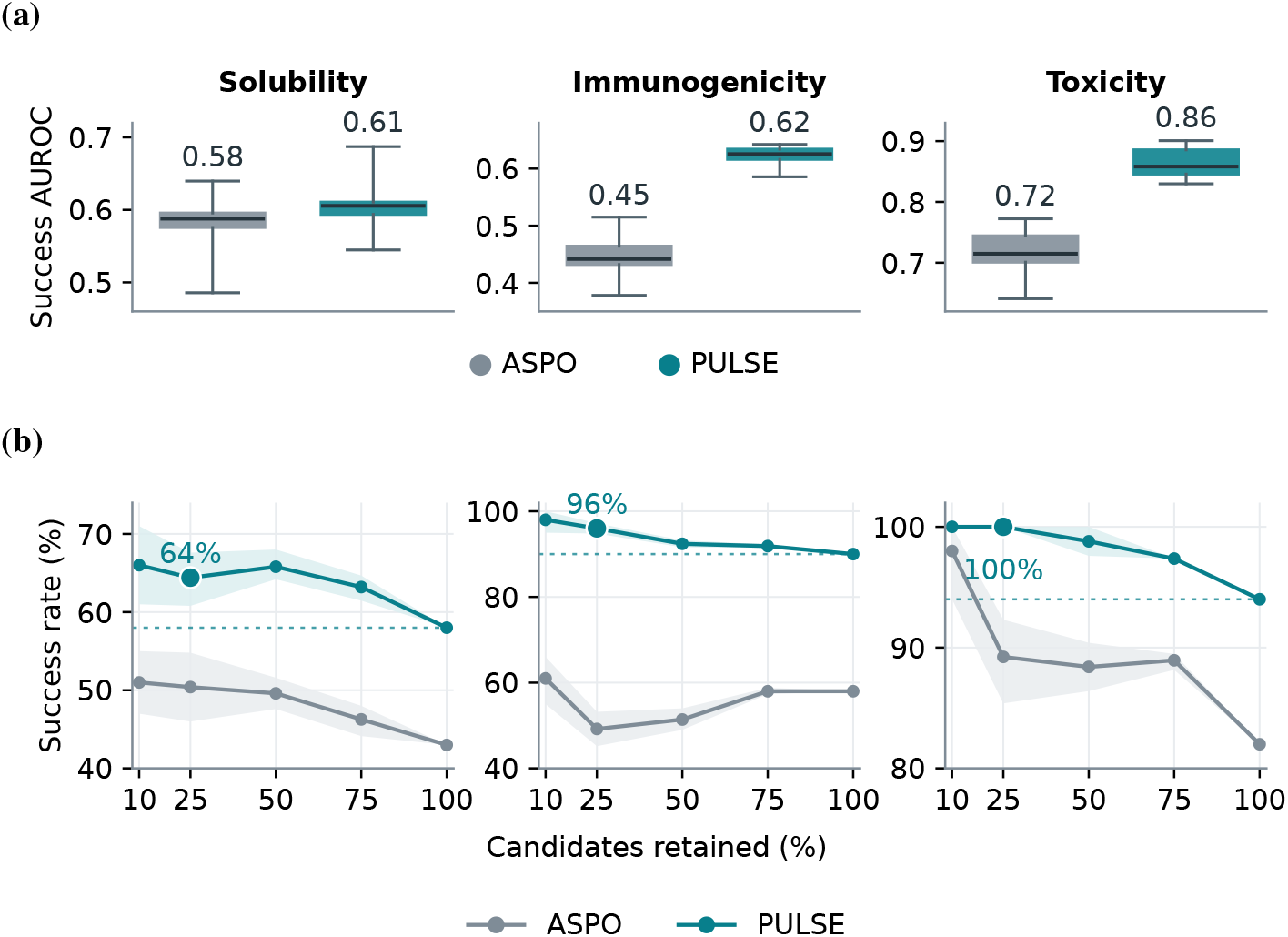
LDS-guided candidate prioritization. (a) Success AUROC across ten reference pools; boxes show median and quartiles, whiskers the range, and labels the mean. (b) Success rate versus LDS-ranked coverage; large points mark the top 25%, dotted lines show overall PULSE success with shading variation across pools for the same candidates.

While LDS provides an effective internal ranking for any upstream steering method, the absolute level of trajectory support determines overall discrimination performance. Because PULSE trajectories achieve systematically higher LDS scores than ASPO across all three properties (Fig. 5), PULSE candidate pools maintain a substantially higher proportion of well-supported trajectories. Consequently, LDS achieves higher AUROC point estimates for PULSE than ASPO across solubility, immunogenicity, and toxicity (Fig. 6a), while delivering higher absolute success enrichment when retaining top-ranked candidates (e.g., the top 25% budget threshold in Fig. 6b).

Crucially, this prioritization framework operates without requiring a trained property oracle at candidate-selection time. While LDS primarily indicates binary trajectory reliability rather than the exact magnitude of property improvement, its method-agnostic ranking fidelity offers a practical, data-efficient strategy for prospective candidate selection under tight experimental budgets.

## 4 Discussion

### 4.1 Global and local trajectory reliability provide complementary information

Trajectory-level reliability can be evaluated across two granularities: **global construction confidence** and **Local Directional Support (LDS)**. Global construction confidence measures trajectory alignment against resampled global steering vectors (Appendix A.3) and LDS evaluates alignment against local, context-matched property directions. These two metrics capture related yet distinct facets of optimization reliability.

Figure 7a shows that global construction confidence reproduced PULSE’s advantage over ASPO for solubility and immunogenicity, demonstrating that population-level vectors remain informative when property directions are globally coherent. However, this global metric broke down for toxicity, where PULSE exhibited lower global confidence than ASPO despite achieving superior optimization outcomes. This failure illustrates that the global confidence can be misleading when property directions exhibit strong background dependence (Appendix B.2).

**Fig. 7:**
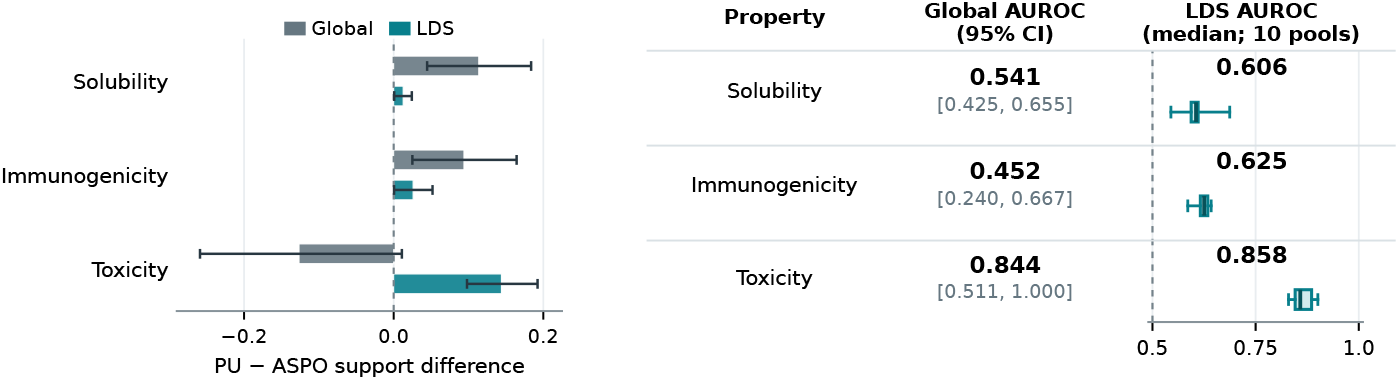
Global and local trajectory reliability. (a) PULSE–ASPO differences in trajectory support; whiskers show 95% paired confidence intervals. (b) Success-ranking AUROC for PULSE candidates. Global values include 95% bootstrap intervals; LDS values and boxplots summarize ten independent reference pools.

By conditioning evaluation on the local contexts encountered along the trajectory, LDS consistently favored PULSE across all three properties, with the largest separation occurring in toxicity (Fig.7a). Rather than replacing global confidence, LDS provides a complementary perspective: global confidence evaluates broad macro-level alignment with the construction population, whereas LDS verifies micro-level validity within immediate representation neighborhoods.

This complementarity translates directly into practical, oracle-free candidate prioritization. While both metrics enable sequence ranking without a trained property predictor, LDS achieved higher AUROC estimates for all three properties (Fig.7b), with the clearest gains in solubility and immunogenicity. For toxicity, the two metrics performed comparably. Global confidence offers a useful population-level baseline, whereas LDS provides essential local fidelity when representation space heterogeneity makes global directions insufficient.

### 4.2 Limitations

In this work, findings are limited to only ESMC model backbone and predictor-based outcomes rather than experimental measurements. LDS requires a suitable independent labeled reference pool, and uses a neighborhood fixed at the starting protein. Its association with optimization success does not establish that local directional alignment causes the improvement. Broader model, optimization budget comparisons and wet-lab validation remain important next steps.

## Acknowledgment

This work has been supported by the Advanced Research Projects Agency for Health (ARPA-H) Award 1AY1AX000053-01.

## Supplementary Information

### A Additional Results

#### A.1 Paired endpoint-score comparison

ASPO and PULSE optimized the same target proteins for each property. Final predicted scores were compared within targets. We estimated 95% confidence intervals for the mean paired difference by resampling target pairs and applied Holm correction to the three paired tests.

PULSE achieved higher mean final predicted scores for all three properties. Each paired difference remained significant after correction (Table 2).

#### A.2 Static steering-vector diagnostics

We assessed global reproducibility by separately bootstrapping the positive and negative construction proteins and rebuilding each steering vector. Let *v*_*m*_ be method *m*’s full-data vector at *l*_select_, and 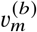 its reconstruction in bootstrap sample *b*. Mean angular deviation, reported in degrees, is

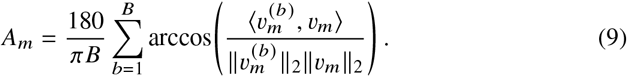

We used *B* = 200; smaller *A*_*m*_ indicates a more reproducible static direction.

Local Static Vector Support (LSVS) measures whether *v*_*m*_ agrees with property-improving directions near a target’s starting background. Each 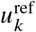 is the unit-normalized positive-minus-negative representation difference of a matched pair from an independent reference pool, computed at *l*_select_. Let 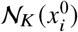 index the *K* = 32 reference pairs whose backgrounds are most similar to the starting sequence 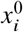 at *l*_pair_. Then

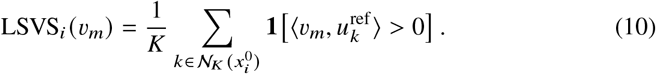

For each target and reference pool, ASPO and PULSE use the same neighborhood. LSVS assesses the static vector rather than the changes realized during optimization.

#### A.3 Global construction confidence

Global construction confidence evaluates realized optimization steps against the boot-strap vectors defined above. For the representation change *δ*_*i,t*_ = *h* (*x*_*i,t* +1_) − *h*(*x*_*i,t*_) made by method *m* at step *t*, we computed

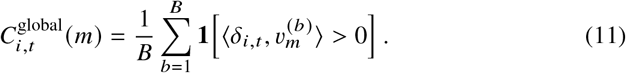

Thus, the score is the fraction of reconstructed global steering directions that agree with the step actually taken. A step with zero representation displacement is assigned a score of 0.5. As for LDS, we summarized each trajectory by the lower quartile of its step scores:

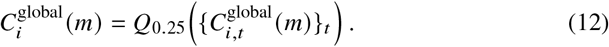

Global construction confidence and LDS evaluate the same realized steps, but use different references: bootstrap vectors from the full construction population for the former, and independent directions selected near the starting protein’s background for the latter.

### B Datasets and Data Cleaning Pipeline

To ensure rigorous evaluation and prevent data leakage, sequence datasets for each property were divided into three mutually exclusive sets:

1. A **main construction set** used to construct ASPO and PULSE steering vectors.
2. A **test set** of target sequences subjected to steering optimization.
3. An **independent LDS reference pool** used exclusively for dynamic trajectory-level reliability evaluation.

#### B.1 Data Cleaning and Leakage Prevention

Prior to evaluation, all duplicate sequences and exact sequence overlaps across the construction set, test set, and LDS reference pool were removed. The LDS reference pools were balanced to contain equal numbers of positive and negative property-improving transitions. Table 4 details the raw candidate counts and final balanced counts.

#### B.2 Background dependence of toxicity-associated directions

We examined whether the toxicity-associated direction changed systematically across protein backgrounds. Among 150 matched pairs, pairwise background similarity was positively associated with pairwise direction similarity (Spearman *ρ* = 0.568; Holm-adjusted permutation *p* = 0.003). Mean direction similarity increased across the background-similarity range (Supplementary Table 3). Toxicity had the largest observed background–direction correlation of the three properties.

### C Hyperparameters and Experimental Configurations

All steering methods were implemented in PyTorch and run with CUDA on an NVIDIA GeForce RTX 5090 GPU. Model seed parameters were initialized with SEED = 42. Layer parameters (*l*_select_, *l*_pair_) and steering intensity *α* were selected per property and maintained across all baseline and ablation comparisons. Table 5 summarizes all hyper-parameter choices.

**Table 5.** Hyperparameter specification across properties and evaluation protocols.

| Category | Hyperparameter | Setting |
|---|---|---|
| Model Backbone | Architecture | ESMC-600M (36 layers, *d* = 1152) |
|  | Pooling Method | Mean pooling over residue positions |
| Solubility | *l*<sub>select</sub>/*l*<sub>pair</sub> | Layer 25 / Layer 35 |
|  | Steering Factor (*α*) | 1.0 |
|  | Test Target Budget (*N*) | 100 sequences |
| Immunogenicity | *l*<sub>select</sub>/*l*<sub>pair</sub> | Layer 35 / Layer 35 |
|  | Steering Factor (*α*) | 2.0 |
|  | Test Target Budget (*N*) | 100 sequences |
| Toxicity | *l*<sub>select</sub>/*l*<sub>pair</sub> | Layer 30 / Layer 30 |
|  | Steering Factor (*α*) | 2.0 |
|  | Test Target Budget (*N*) | 50 sequences |
| Evaluation Diagnostics | Local Neighborhood Size (*K*) | 32 |
|  | Trajectory LDS Summary | *Q*<sub>0.25</sub> |
|  | Random Pairing Seeds | 5 seeds (17, 29, 43, 71, 101) |

The oracle property scores *f*(·) were obtained from the evaluation predictors used in [15]. For solubility, we used the binary fitness predictor described by Huang et al. [7]: an ESM-2-650M feature extractor with a prediction head trained on the pre-processed DeepSol data, comprising 28,972 soluble and 40,448 insoluble proteins [8]. For immunogenicity and toxicity, we used property-specific fine-tuned VenusVaccine predictors. Their dual-attention backbone integrates pretrained protein sequence and structure representations [10]; the corresponding labeled data sources were ImmunoDB for immunogenicity [10] and ToxDL 2.0 for toxicity [23]. These predictors scored both starting and optimized sequences to evaluate property changes. They were not used to rank candidates by LDS.

## References

1. Biswas, S., et al.: Low-N protein engineering with data-efficient deep learning. Nat. Methods 18, 389–396 (2021). doi:10.1038/s41592-021-01100-y

2. Chu, H.Y., et al.: Accurate top protein variant discovery via low-N pick-and-validate machine learning. Cell Syst. 15, 193–203.e6 (2024). doi:10.1016/j.cels.2024.01.002

3. Efron, B., Tibshirani, R.J.: An Introduction to the Bootstrap. Chapman & Hall, New York (1993)

4. ESM Team: ESM Cambrian: revealing the mysteries of proteins with unsupervised learning. Biohub (2024).

5. Hie, B.L., et al.: Efficient evolution of human antibodies from general protein language models. Nat. Biotechnol. 42, 275–283 (2024). doi:10.1038/s41587-023-01763-2

6. Hayes, T., et al.: Simulating 500 million years of evolution with a language model. Science 387, 850–858 (2025). doi:10.1126/science.ads0018

7. Huang, L.-K., Zhu, R., He, B., Yao, J.: Steering protein language models. In: Proceedings of the 42nd International Conference on Machine Learning. PMLR 267, 26247–26260 (2025).

8. Khurana, S., et al.: DeepSol: a deep learning framework for sequence-based protein solubility prediction. Bioinformatics 34, 2605–2613 (2018). doi:10.1093/bioinformatics/bty166

9. Kuhn, H.W.: The Hungarian method for the assignment problem. Nav. Res. Logist. Q. 2, 83–97 (1955). doi:10.1002/nav.3800020109

10. Li, S., Tan, Y., Ke, S., Hong, L., Zhou, B.: Immunogenicity prediction with dual attention enables vaccine target selection. In: International Conference on Learning Representations (2025).

11. Lin, Z., et al.: Evolutionary-scale prediction of atomic-level protein structure with a language model. Science 379, 1123–1130 (2023). doi:10.1126/science.ade2574

12. Luo, Y., et al.: ECNet is an evolutionary context-integrated deep learning framework for protein engineering. Nat. Commun. 12, 5743 (2021). doi:10.1038/s41467-021-25976-8

13. Madani, A., et al.: Large language models generate functional protein sequences across diverse families. Nat. Biotechnol. 41, 1099–1106 (2023). doi:10.1038/s41587-022-01618-2

14. Meier, J., Rao, R., Verkuil, R., Liu, J., Sercu, T., Rives, A.: Language models enable zero-shot prediction of the effects of mutations on protein function. In: Advances in Neural Information Processing Systems 34 (2021).

15. Qayyum, A.B.A., Zhou, Y., Qian, X., Yoon, B.-J.: Optimizing protein design through uncertainty-weighted steering of protein language models. Bioinformatics 42(Suppl. 2), btag436 (2026). doi:10.1093/bioinformatics/btag436

16. Qian, J., et al.: Accelerating protein design by scaling experimental characterization. Nat. Commun. 17, 9945 (2026). doi:10.1038/s41467-026-76740-9

17. Rimsky, N., Gabrieli, N., Schulz, J., Tong, M., Hubinger, E., Turner, A.: Steering Llama 2 via contrastive activation addition. In: Proceedings of ACL, pp. 15504–15522 (2024).

18. Rives, A., et al.: Biological structure and function emerge from scaling unsupervised learning to 250 million protein sequences. Proc. Natl. Acad. Sci. USA 118, e2016239118 (2021). doi:10.1073/pnas.2016239118

19. Sarkisyan, K.S., et al.: Local fitness landscape of the green fluorescent protein. Nature 533, 397–401 (2016). doi:10.1038/nature17995

20. Starr, T.N., Thornton, J.W.: Epistasis in protein evolution. Protein Sci. 25, 1204–1218 (2016). doi:10.1002/pro.2897

21. Turner, A.M., et al.: Activation addition: steering language models without optimization. 2308.10248 (2024). doi:10.48550/arXiv.2308.10248

22. Weinreich, D.M., Delaney, N.F., DePristo, M.A., Hartl, D.L.: Darwinian evolution can follow only very few mutational paths to fitter proteins. Science 312, 111–114 (2006). doi:10.1126/science.1123539

23. Zhu, L., et al.: ToxDL 2.0: protein toxicity prediction using a pretrained language model and graph neural networks. Comput. Struct. Biotechnol. J. 27, 1538–1549 (2025). doi:10.1016/j.csbj.2025.04.002

